# Extracellular spike waveform dissociates four functionally distinct cell classes in primate cortex

**DOI:** 10.1101/537738

**Authors:** Caterina Trainito, Constantin von Nicolai, Earl K. Miller, Markus Siegel

## Abstract

Understanding the function of different neuronal cell types is key to understanding brain function. However, cell type diversity is typically overlooked in electrophysiological studies in awake behaving animals. Here, we show that four functionally distinct cell classes can be robustly identified from extracellular recordings in several cortical regions of awake behaving monkeys. We recorded extracellular spiking activity from dorsolateral prefrontal cortex (dlPFC), the frontal eye field (FEF), and the lateral intraparietal area of macaque monkeys during a visuomotor decision-making task. We employed unsupervised clustering of spike waveforms, which robustly dissociated four distinct cell classes across all three brain regions. The four cell classes were functionally distinct. They showed different baseline firing statistics, visual response dynamics, and coding of visual information. While cell class-specific baseline statistics were consistent across brain regions, response dynamics and information coding were regionally specific. Our results identify four waveform-based cell classes in primate cortex. This opens a new window to dissect and study the cell-type specific function of cortical circuits.

## Introduction

Neuronal cell types are central to brain function. The unique physiology, morphology and connectivity of different cortical interneurons and pyramidal cells shapes their functional role in local and large-scale circuit operations (Tremblay, Lee, and Rudy 2016; Lee, Koch, and Mihalas 2017; Zeng and Sanes 2017). Cell-type specific neuronal properties shape characteristic circuit oscillations associated with various computational and cognitive processes (Womelsdorf et al. 2014; Veit et al. 2017; Siegel, Donner, and Engel 2012) Thus, knowledge about cell types and their role in cortical circuits is key to understanding brain function.

The assessment of cell types ideally relies on morphological, molecular or genetic markers (The Petilla Interneuron Nomenclature Group (PING) et al. 2008; Tasic et al. 2018). While these markers are often not available for extracellular electrophysiology studies, also firing patterns and action-potential shape provide some handle on cell-type diversity. *In vitro* studies first demonstrated that morphologically identified pyramidal cells and GABA-ergic interneurons differ in firing patterns and action potential shape. Pyramidal cells show regular, low-rate firing patterns and have broad spike waveforms (‘broad-spiking’ units), while inhibitory cells fire at sustained high frequencies with characteristically thin spike waveforms (‘narrow-spiking’ units) (Connors and Gutnick 1990; McCormick et al. 1985; González-Burgos et al. 2004). These intracellular features map on to extracellular features recorded *in vivo* (Nowak et al. 2003).

Based on these findings, several studies have inferred putative cell types from extracellular single-unit activity. In primate prefrontal cortex (Constantinidis and Goldman-Rakic 2002; Diester and Nieder 2008; Johnston, DeSouza, and Everling 2009; Ardid et al. 2015), FEF (Cohen et al. 2009; Thiele et al. 2016), IT (Tamura et al. 2004; Mruczek and Sheinberg 2012) and V4 (Mitchell, Sundberg, and Reynolds 2007; Vinck et al. 2013), spike waveform width is bimodally distributed, indicative of the known separation between excitatory cells and inhibitory interneurons. The proportion of narrow-spiking units in these studies (around 15-25%) is consistent with anatomical estimates of the proportion of GABAergic cells in the cortex (Hendry et al. 1987; note laminar variability: Rudy et al. 2011; Gabbott and Bacon 1996). Firing properties, selectivity and task-related modulations differ between broad- and narrow-spiking units, further supporting the physiological interpretation of distinct cell types (Katai et al. 2010; Ardid et al. 2015). In sum, so far waveform width provides a limited handle on cell-type diversity in the primate brain, allowing to dissociate two broad classes of putative cell types (excitatory vs. inhibitory). However, in order to better understand cell-type specific mechanisms and functions more cell types need to be identified. Furthermore, cell-type classification and functions need to be compared across different cortical regions.

To address this, we characterized putative cortical cell types based on spike waveforms in a large dataset of extracellular recordings from three different cortical regions (FEF, dorsolateral PFC and LIP) in two macaque monkeys (Siegel, Buschman, and Miller 2015). In contrast to the typically reported dichotomy between broad-spiking and narrow-spiking units, we were able to distinguish four cell classes based on waveform shape. These four distinct cell classes were confirmed by cell-class specific firing patterns, response dynamics, and information coding. While the four cell-classes were consistently found across all cortical regions, their functional profiles differed between areas. These findings open a new window into cell-type specific functions in awake behaving animals.

## Results

### Cell class separation based on spike waveform

We analyzed data from 2488 single-units recorded in FEF (793), dlPFC (1050) and LIP (645) of two macaque monkeys (Fig. 1). In a first step, we identified different cell classes in a purely data-driven fashion based on spike waveform. To increase statistical power, we pooled the data across all cortical regions and, for each unit, quantified two parameters of the spike waveform that contribute to the overall spike-width: trough-to-peak duration and repolarization-time (Figure 1A). Trough-to-peak duration is the interval between the global minimum of the curve and the following local maximum. Repolarization time is the interval between the late positive peak and the inflection point of the following falling flank of the curve. Although correlated, these two measures capture different aspects of the intracellular action potential — the speed of depolarization and of the subsequent after-hyperpolarization (Henze et al. 2000) — that are both distinguishing features of neuronal cell types (Nowak et al. 2003). All 2488 waveforms were scored on the two measures to obtain a two-dimensional feature space for classification (Fig. 1B).

**Figure 1.**
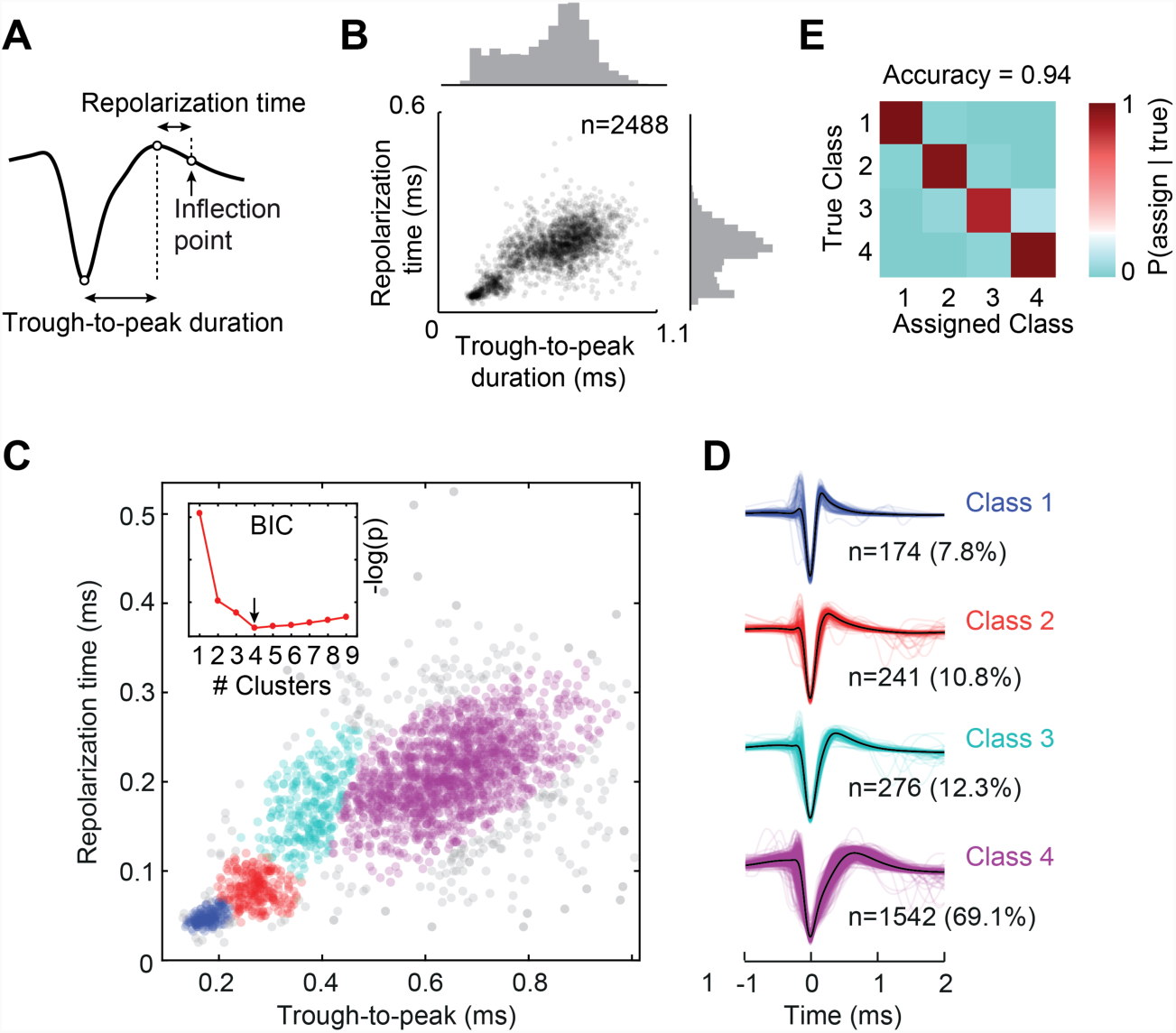
Cluster analysis of extracellular spike waveforms. (A) Illustration of the two spike waveform features used for classification. (B) 2D feature space and marginal distributions of waveforms for all recorded single-units. (C) Clusters of spike waveforms obtained from the Gaussian mixture (GM) model. Single-units are assigned to the cluster with the highest posterior probability. Grey data points are excluded outliers: an initial fifth high-variance cluster and outliers of the GM distribution (n=281). The inset shows the negative log-likelihood of the BIC as a function of the number of clusters after outlier removal (D) All waveforms by cell class (average waveforms in black). (E) Class separation. To quantify the separation of the four clusters, 10^4^ data points were randomly generated from the fitted GM distribution, and their true cluster was compared with their assigned cluster. The classification outcome is shown by the confusion matrix of marginal probabilities. Accuracy is the mean of the four diagonal probabilities.

To identify different cell classes in an unsupervised way, we performed a two-dimensional cluster analysis of the waveform parameters (Gaussian mixture model). We used the Bayesian information criterion (BIC) to select the number of Gaussian components in the model. The BIC showed a global minimum for four components indicating four distinct waveform classes (Fig. 1C) Ranging from narrow to wide waveforms, the four classes comprised 7.8%, 10.8%, 12.3% and 69.1% of the sample, respectively. Thus, most units were attributed to the widest waveform class (class 4). We quantified cluster separation by calculating the probability to correctly classify each cell class based on the Gaussian mixed model underlying the clustering (Fig. 1D). The average classification accuracy across all four classes was 94%, indicating well-separated clusters.

### Cell classes across cortical regions

We next investigated if the waveform-based cell classes were robust across different cortical areas (Figure 2). Splitting the data by areas revealed that the four classes were unequally distributed across cortical regions (χ^2^ omnibus test, p<0.001; Figure 2A). Thus, we investigated if the four waveform clusters were consistently identified within each region. Indeed, clustering run separately on each area consistently returned four classes with the same overall structure (Figure 2B).

**Figure 2.**
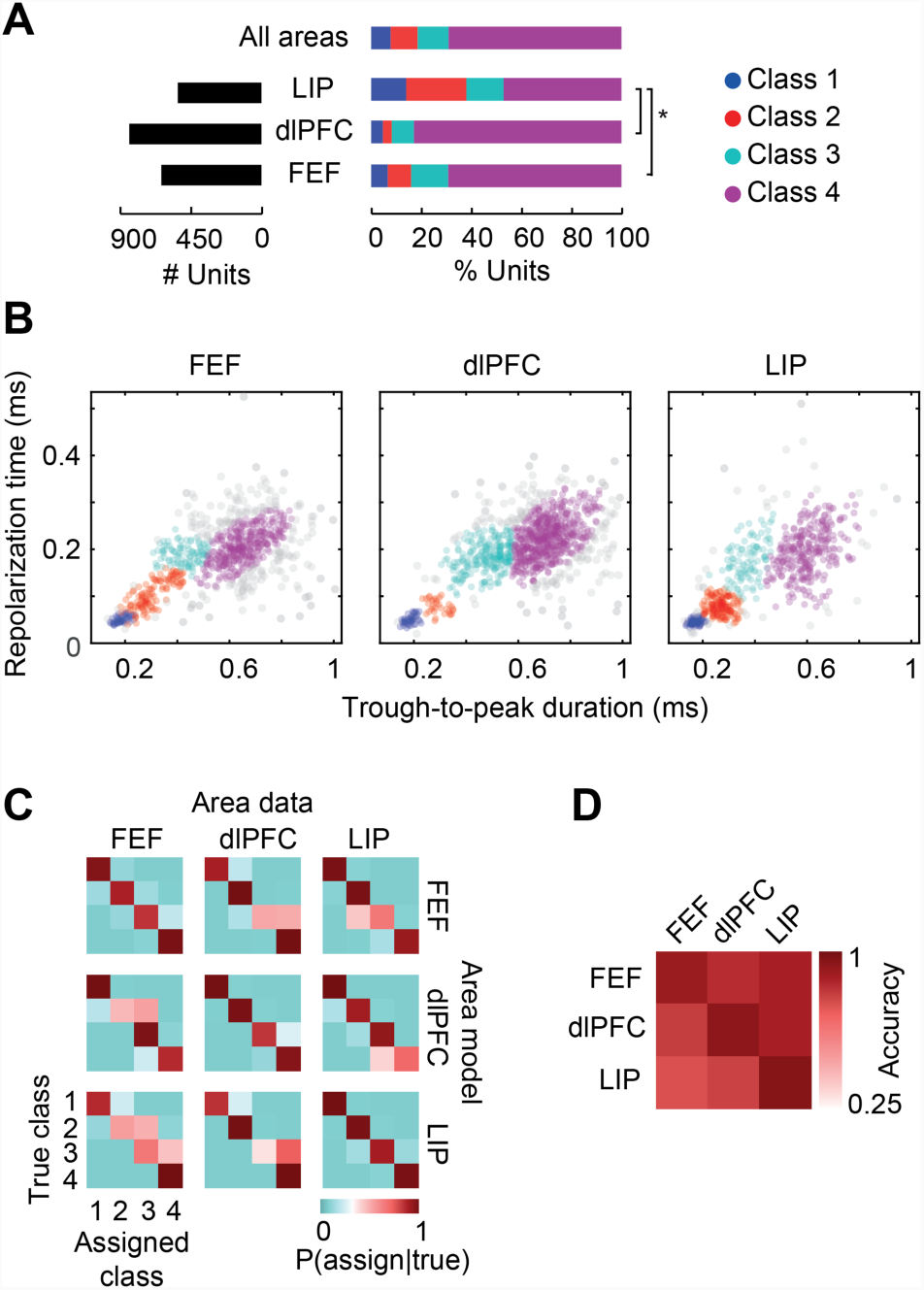
Reliability of waveform clustering across cortical regions. (A) Distribution of units across cortical regions and cell classes. Brackets indicate significant post-hoc χ^2^ tests for different cell-class distributions across areas (p<0.05, Bonferroni-corrected. (B) Waveform feature spaces with clustering run separately for FEF, dlPFC and LIP. (C) Cluster separation and similarity of cell classes across areas. Confusion matrices on the diagonal show separation of the four clusters for each area’s own GM model (as in Figure 1E). For all other area pairs, confusion matrices measure the similarity between the same cell class clusters in the two areas. Cluster similarity is estimated by randomly generating 10^4^ data points from one area’s GM distribution (“area data”) and classifying them based on the GM distribution of the other area (“area model”). (D) Mean diagonal probabilities of confusion matrices in (C).

To estimate the cluster separability within each area, we quantified the probability to correctly classify each class based on the Gaussian mixed model within each region (Figure 2C diagonal panels). Furthermore, to estimate the waveform class similarity across brain regions, we quantified cross-classification accuracy between different regions, i.e. we trained and tested the classifier on different regions (Figure 2C off-diagonal panels). For both cases and across all brain regions, classification accuracy was above 75% (Figure 2D). This indicates both, a consistently high separation between the four clusters within each region and a high overlap of each clusters across regions. In sum, the four waveform based cell classes were robustly and similarly observed across the three cortical regions.

### Firing statistics of cell classes

What are the functional properties of the four putative identified cell types? If the four spike waveform clusters reflect distinct physiological cell types, the corresponding units should show different functional characteristics. We started by examining firing statistics during the blank fixation baseline of a flexible visual decision-making task (Figure 3). For each neuron, we computed four statistics: mean firing rate across trials (FR), Fano factor (variance over mean of spike counts across trials, FF), coefficient of variation of the inter-spike interval distribution (CV_ISI_) and burst index (BI). Both, Fano factor and CV_ISI_ are mean-standardized measures of dispersion that reflect firing variability, with an expected value of 1 for Poisson firing and values below 1 indicating more regular firing (Shinomoto, Shima, and Tanji 2003). Burst index was defined as the ratio between the observed proportion of bursts (inter-spike intervals <5 ms) and the proportion of bursts expected for a Poisson process with equal mean rate (Constantinidis and Goldman-Rakic 2002). One-way ANOVAs showed significant class separation on all four measures (all p<0.05) (Figure 3A). Firing rate was highest for class 1 units (narrow waveforms), followed by the two intermediate-waveform classes 2 and 3 (not significantly different from each other), and lowest for class 4 (broad-spiking units). Fano factor showed a similar pattern: Class 4 had the lowest Fano factor and therefore more regular firing, also confirmed by the low CV_ISI_. These results agree with the classical designation of narrow-waveform neurons as fast-spiking (FS) and broad-waveform neurons as regular-spiking (RS) (McCormick et al. 1985; Connors and Gutnick 1990; Nowak et al. 2003). On the other hand, the intermediate-waveform Class 3 was more likely to fire in bursts than any other class.

**Figure 3.**
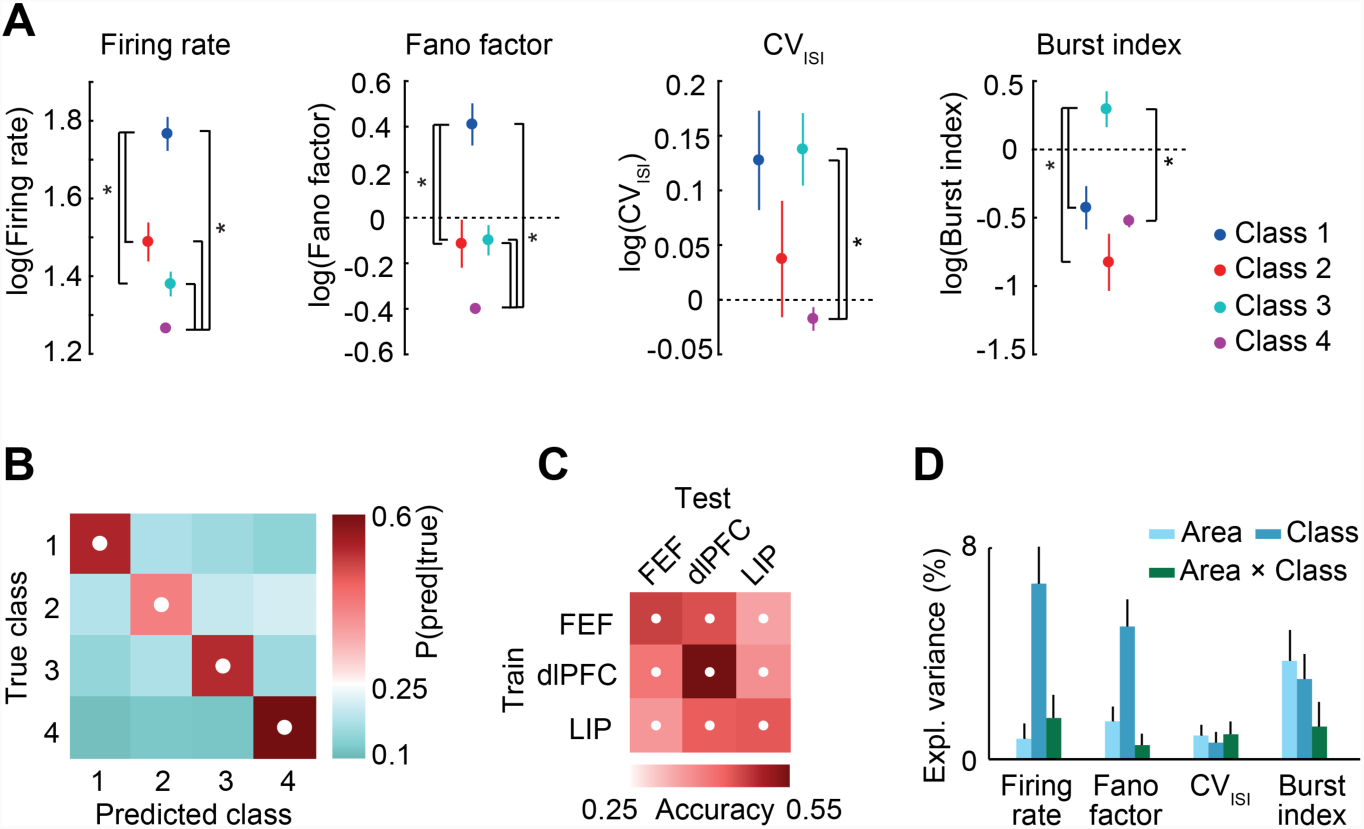
Cell-class specific baseline firing statistics. (A) Firing statistics by cell class. All measures were computed during a 500 ms baseline fixation period of the trial. CV_ISI_: coefficient of variation of the inter-spike interval distribution. Burst index: proportion of inter-spike intervals <5 ms over the proportion expected from a Poisson neuron. Brackets indicate significant post-hoc pairwise differences (p<0.05, FDR-corrected). (B) Confusion matrix for supervised classification of cell classes using the four baseline firing statistics as features. White dots indicate overall performance significantly above chance (binomial test, p<0.05, FDR-corrected). (C) Mean diagonal probabilities (“accuracy”) for cross-area classification. Classifiers trained on data from one area (“Train“) were used to predict class labels of the other area (“Test“). White dots indicate significant class prediction (permutation test, p<0.05, FDR-corrected). (D) Variance explained by cell class, cortical area and their interaction in a two-way ANOVA performed on each firing statistic by area. All main and interaction effects were significant (p<0.05).

### Firing statistics validate four cell classes

The significant differences of firing statistics between cell classes support the conclusion that the four waveform-defined cell classes reflect distinct physiological cell types. To further validate this conclusion, we employed a machine-learning approach: assuming the waveform-based classes as ground truth, we trained a multivariate classifier (SVM, support vector machine) to decode these four cell classes from all four firing statistics. Again, if the four waveform clusters reflect distinct cell types, class membership should be predictable from functional cell properties. Indeed, we were able to significantly predict all four cell classes with high classification accuracy (Figure 3B; class. accuracy: 0.53±0.02; mean±SD over 50 area-stratified subsamples,all p<0.05, FDR-corrected, binomial test).

The classification approach allowed us to test if class-specific functional properties were stable across cortical regions. We trained classifiers on data from one cortical area and tested them on data from a different area (cross classification). Classification performance always exceeded chance level (permutation test, all p<0.05, FDR-corrected; Figure 3C) suggesting that cell classes maintain their functional profiles across regions. This was also indicated by the univariate measures broken down by area (Supplementary Figure 1). While there was a significant interaction between cell class and area (two-way ANOVAs, all measures p<0.05), the cell-class effect was significant for all statistics, and for firing rate and Fano factor, most of the variance was independently explained by cell class (Figure 3D). In sum, firing statistics differed between, and thus validated, four waveform-based cell classes that were robust across cortical regions.

### Cell class-specific firing dynamics

Next, we investigated if the four cell classes differed in their firing dynamics in response to a sensory stimulus (Figure 4). We characterized each neuron’s response to presentation of a visual cue that indicated the upcoming task condition for each trial of the perceptual decision-making task (see schematic of behavioral paradigm in Figure 4A). We computed peristimulus time histograms (PSTH) in a window including the baseline fixation period (0.5 s) and the subsequent cue period (1 s).

**Figure 4.**
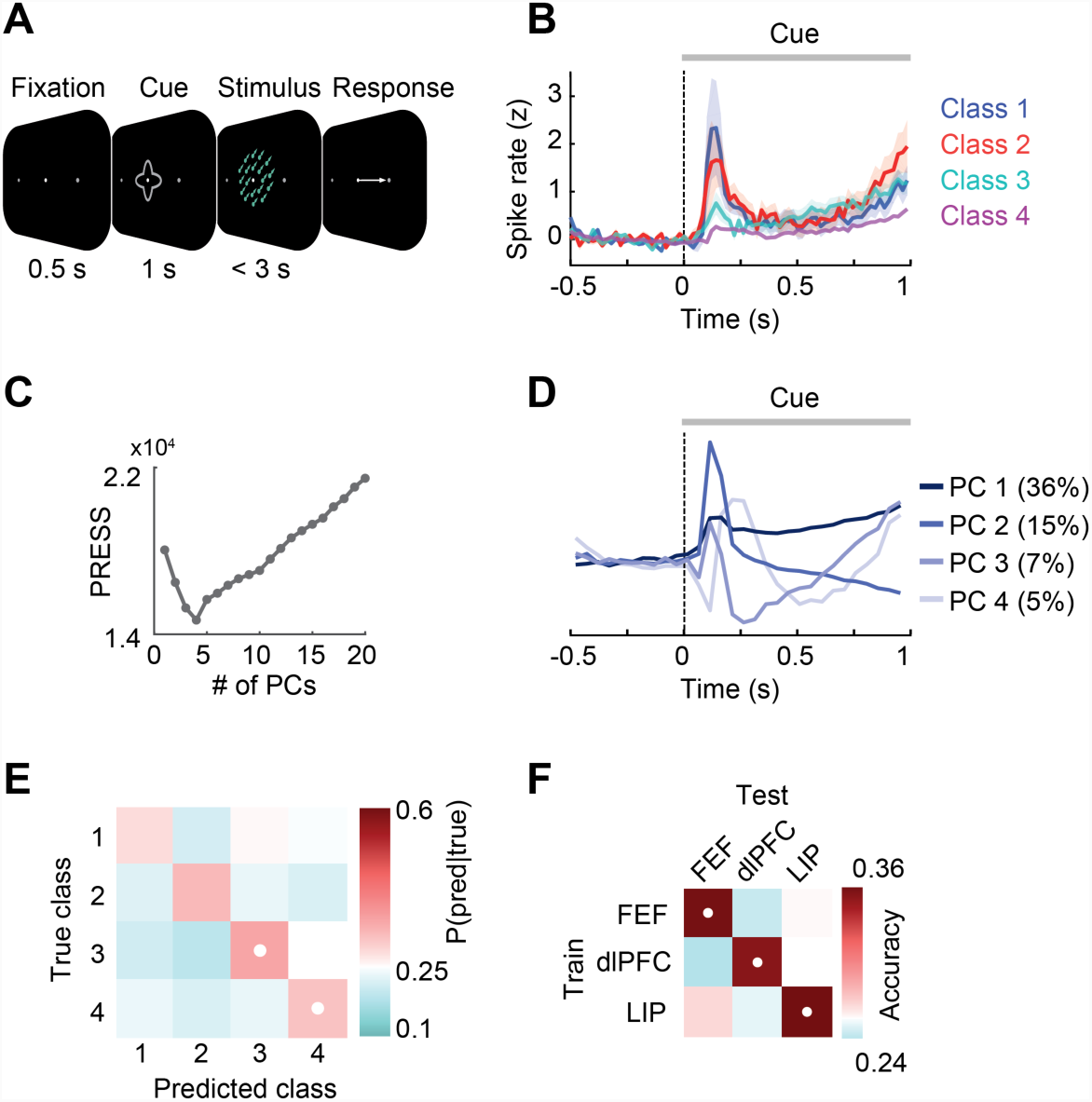
Cell-class specific response dynamics. (A) Schematic of the behavioral task. (B) Average PSTH for each cell class. PSTHs were z-scored on the mean and SD of the baseline period across trials. PSTH means and SD (error bars) were calculated after stratifying cell-class proportions across areas. (C) Selection of number of PCA components via cross-validation. Reconstruction error (PRESS, Prediction Residual Error Square Sum) as a function of number of principal components. (D) Significant principal components (PCs). Percentages denote the variance explained by each PC. (E) Confusion matrix for supervised classification of cell classes using PCA projections of the PSTHs. White dots indicate significant class prediction (binomial test, p<0.05, FDR-corrected). (F) Mean diagonal probabilities (“accuracy”) for cross-area classification. Classifiers trained on data from one area (“Train“) were used to predict class labels of the other area (“Test“). The PCA transformation was estimated on the training area and applied to data of the test area. White dots indicate significant class prediction (permutation test, p<0.05, FDR-corrected).

The cell-class specific PSTHs pooled across regions suggested differences of the response dynamics between cell classes (Figure 4B). To statistically assess this, we captured firing dynamics in a low-dimensional space. We performed a Principal Component Analysis (PCA) of the PSTHs of all neurons pooled across regions. We then used a cross-validation procedure to estimate the effective rank of the dataset, i.e. the number of underlying orthogonal dynamical features or principle components. We found four significant components (Figure 4C), explaining 63% of the response-dynamics’ variance. Visual inspection of the components (Figure 4D) showed that most variance was captured by dynamical components reflecting both tonic, ramp-like firing (PC1) and time-localized evoked responses (PC2) including slower modulations (PC3, PC4).

We projected each neurons PSTH on the four significant PCs and used the resulting low-dimensional representation of the response dynamics as features for multivariate decoding. Importantly, we normalized PSTHs on the mean spike rate of the baseline period. This ensured that the decoder was not classifying merely based on differences in overall activity levels, but that it specifically probed cue-related responses. Furthermore, to rule out a confound of the region-specific distribution of cell classes, we stratified the number of cells per class across regions (see Methods). We found that cell classes 3 and 4 could be significantly decoded confirming cell class-specific PSTH dynamics (Figure 4E; binomial test on confusion matrix probabilities, all p<0.05, FDR-corrected for multiple comparisons; classifier accuracy: 0.32±0.03; mean±SD).

Sensory response latencies and dynamics differ between brain regions. Thus, we hypothesized that cell-class specific response dynamics would be area specific. Indeed, we found that, unlike for baseline firing statistics, cell class decoding based on response dynamics did not significantly generalize across areas (Figure 4F). Accordingly, the cell classes’ response dynamics showed dissimilar patterns in the three areas (Supplementary Figure 2).

### Cell class-specific information coding

If cell classes vary in their cue-evoked response dynamics, do they also differentially code for specific cues? To address this question, for each neuron, we quantified the amount of cue information encoded by its firing rate, by measuring the amount of firing rate variance across trials explained by cue identity (Figure 5; ANOVA, 4 cues). We then trained a classifier to decode cell classes based on cue information. Again, we controlled for a region confound by stratifying cell classes across areas. Furthermore, to control for confounds due to firing statistics, before classification, we regressed out linear dependencies of cue information on baseline firing statics (firing rate, Fano factor, coefficient of variation of the ISI distribution, burst index). We found that cell classes 2 and 4 could be significantly decoded from cue information (Figure 5B). Furthermore, we found that the cell class specificity of cue information was region-specific. Cross-area classification performance was low (Figure 5C) and the pattern of cue information across cell classes differed between regions (Supplementary Figure 3). Thus, while neurons of class 2 and 4 on the whole carried different cue-related information, the direction of the effect was area specific.

**Figure 5.**
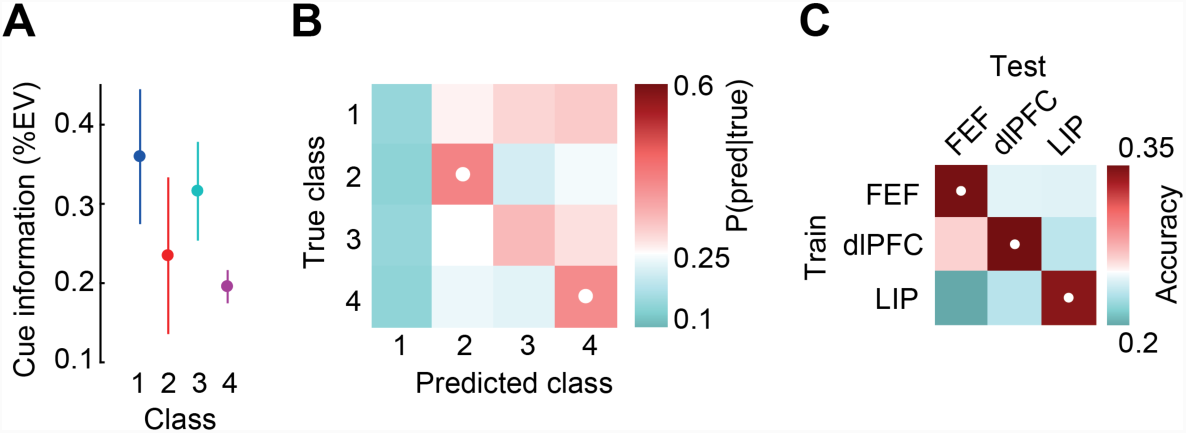
Decoding of cell classes from cue-related information. (A) Cue information by cell class. Cue information was quantified as spike rate variance (ω^2^) in the late cue period (500-1000 ms from stimulus onset) across trials explained by cue identity. The four baseline firing statistics (firing rate, Fano factor, CV_ISI_, burst index) were regressed out. Average information was calculated after stratifying cell-class proportions across areas. Cue information significantly differed between cell classes (one-way ANOVA, p<0.05). (B) Confusion matrix for supervised classification of cell classes from cue information. White dots indicate significant classification performance (binomial test, p<0.05, FDR-corrected). (C) Mean diagonal probabilities (“accuracy“) for cross-area classification using cue information. Classifiers trained on data from one area (“Train“) were used to predict class labels of the other area (“Test“). White dots indicate significant classification (permutation test, p<0.05, FDR-corrected).

### Specificity of functional properties

Having established that the four cell classes differ in baseline activity, response dynamics and information coding, we pooled together all three feature sets to construct an ‘omnibus’ decoder to predict cell class (Figure 6). To assess each feature’s relative contribution to classification, we recast the problem in a linear framework (LDA) and used the univariate class effects, normalized to a common scale, as a proxy for feature importance. We computed feature importance for each of the six pair-wise cell classifications (Figure 6C) and then averaged to show the overall weightings (Figure 6D). Furthermore, we compared cell-class classification accuracy (Figure 6E) and area specificity (Figure 6F) for each individual feature set and all combined sets.

**Figure 6.**
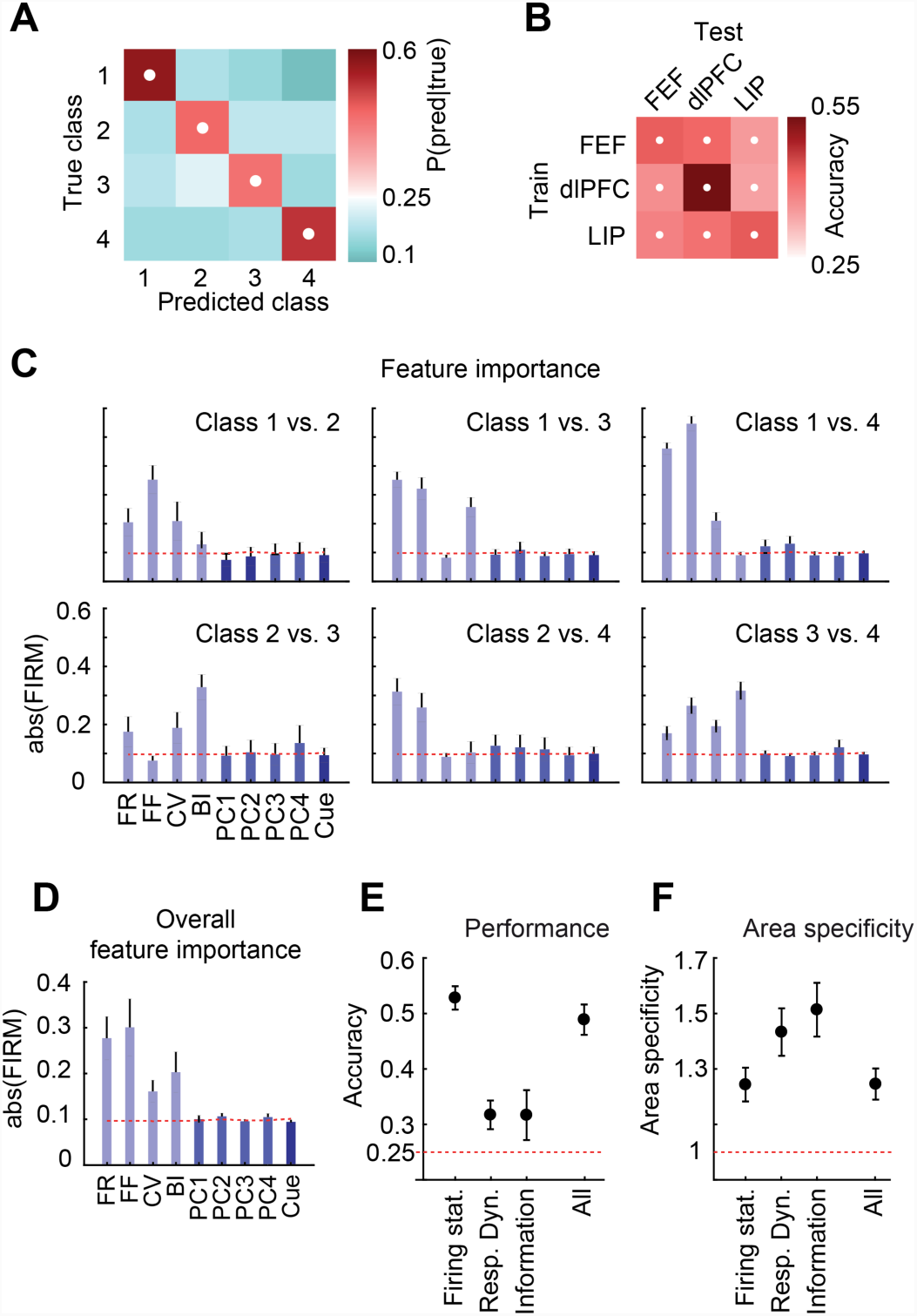
Decoding of cell classes from all combined functional measures. (A) Confusion matrix for supervised classification of cell classes using all functional measures as features (four baseline firing statistics, PCA projections of PSTH, cue information). White dots indicate significant classification (binomial test, p<0.05, FDR-corrected). (B) Mean diagonal probabilities for cross-area classification. For the PSTH features, the PCA transformation was estimated on the training area and applied to data of the test area. White dots indicate significance (permutation test, p<0.05, FDR-corrected). (C) Feature importance for all features derived from pairwise linear classifiers quantified as the magnitude of FIRM (Feature Importance Ranking Measure). Error bars are SD across 50 area-stratified subsampled datasets. The red line shows reference FIRM values for a ‘null’ classifier with shuffled class labels (FR, mean firing rate; FF, Fano factor; CV, coefficient of variation of the ISI distribution; BI, burst index; PC1–PC4, PSTH projections on the first four PCs; Cue: Cue information. (D) Feature importance for all features, averaged across the six pairwise binary classifiers. Error bars show the SEM across the binary classifiers. The red line shows reference FIRM values for ‘null’ classifiers using shuffled class labels. (E) Accuracy across cell classes for all four classifiers. Accuracy is the mean diagonal probability of the confusion matrix. Error bars show the SD across 50 area-stratified datasets. The red dashed line indicates chance-level accuracy of 0.25. (F) Area specificity for all four classifiers. Area specificity is computed as the ratio between average within-area and cross-area classification accuracy. The red dashed line indicates the value expected for perfect generalizability across areas. Error bars show the SD across 50 area-stratified datasets.

These analyses showed that cell classes were most strongly separable by the four baseline-firing statistics. This separation was most consistent across cortical regions (compare Figure 3C), suggesting that cell types maintain their basic firing properties even when embedded in functionally diverse areas. While also showing class effects, cue-related response dynamics and information coding were less cell-class specific, and to a greater extent reflected area-specific process. Furthermore, pairwise feature importance (Figure 6C) showed that cell-class separation differed for distinct response dynamics depending on which two classes were being compared.

Finally, to rule out that the above results merely reflected different spike waveforms or functional cell properties for the two monkeys rather than distinct cell classes, we independently repeated the cell-class decoding for each of the two animals using all functional measures as features (four baseline firing statistics, PCA projections of PSTH, cue information). This revealed very similar independent results for both animals (Figure 7).

## Discussion

We employed a large dataset of electrophysiological recordings in awake, behaving monkeys to distinguish cortical cell types based on extracellular spike waveform. Across dlPFC, FEF and LIP, we robustly identified four distinct cell types that showed distinct functional properties in terms of baseline firing statistics, sensory response dynamics, and information coding.

### Four waveform-based classes

Our results go beyond previous studies that dissociated only two cell classes (narrow-spiking putative interneurons vs. broad-spiking putative pyramidal cells) based on extracellular spike waveform in monkeys (Constantinidis and Goldman-Rakic 2002; Mitchell, Sundberg, and Reynolds 2007; Diester and Nieder 2008; Oemisch et al. 2015; Thiele et al. 2016; Mruczek and Sheinberg 2012; Johnston, DeSouza, and Everling 2009; Vinck et al. 2013; Hussar and Pasternak 2009). An important factor for this advance is likely that we employed a two-dimensional feature space for waveform classification. We considered two highly informative waveform measures that have a known physiological relationship to cell type-specific action potential dynamics (Nowak et al. 2003). Most previous work using peak-to-trough duration as single waveform feature found a clear bimodal distribution, which justified a two-class scheme. In our data, only the time for repolarization measure showed two distinct modes (see marginal histogram in Figure 1B), while peak-to-trough duration likely consisted of several latent components. Together, these measures allowed for defining four bivariate clusters that were less discriminable when projected only onto one dimension (see also Barthó et al. 2004; Insel and Barnes 2015).

Importantly, owing to the high statistical power of our large dataset, we were able to use unsupervised methods to discover patterns in the data. We performed classification without *a priori* definition of the number of clusters. We also determined class assignments by purely statistical criteria, instead of using pre-specified thresholds (i.e. specific values of spike width). This avoided potential confounds due to *a priori* parameter selection.

A cross-classification analysis revealed that waveform clustering was robust across cortical regions. This has two important implications, First, while increasing statistical power, pooling of single-units across FEF, dlPFC and LIP meant that clustering outcomes could be biased by cortical area. For example, if there were only two true classes that occupied slightly different regions of the 2D feature space depending on the recording area, then the whole sample would spuriously appear to contain multiple latent classes. This was not the case, as we ascertained by re-running the unsupervised cluster analysis independently on data from the three areas which reliably revealed four cell classes with comparable statistical structure in each area (Figure 2). Second, this finding supports the notion of cell types as stable physiological entities serving specific roles at the level of cortical microcircuits and columns, rather than at the level of cortical regions (Harris and Shepherd 2015). However, it should be noted that research on area specificity of cell types is still in its infancy (Zeng and Sanes 2017) and that excitatory cells indeed show distinct transcription profiles across cortical regions (Tasic et al. 2018).

### Waveform-width

Our results add to a growing body of evidence suggesting action-potential width as a versatile cell-class marker. In monkey dlPFC *in vitro*, a morphologically-confirmed ‘adapting non-pyramidal’ cell class shows a distinct intermediate spike waveform, significantly different in width from that of both regular-spiking and fast-spiking cells (González-Burgos et al. 2004). Among twelve intracellularly measured physiological parameters, action potential duration had the largest effect size (Krimer et al. 2005). The discriminating power of spike width has been systematically tested in an analysis of electrophysiologically-defined cell types (‘e-types’) in rat S1 (Druckmann et al. 2013). Here, spike width was ranked as best-discriminating feature out of 38 electrophysiological measures. Taken together, these and our present results suggest that spike waveform is a sufficiently sensitive and specific marker to dissociate more than two cell classes from extracellular recordings.

### Functional dissociation between cell classes

We found significant differences of functional properties between waveform based cell classes, in terms of firing statistics, response dynamics and information coding. For the present data, no ground truth on cell class membership was available. Thus, functional differences provide an important independent validation of the waveform based cell classes. In accordance with the distinct functional roles of FEF, dlPFC and LIP, cell-class specific response dynamics and information coding varied substantially across areas (Siegel, Buschman, and Miller 2015). In contrast, baseline firing-statistics were consistently cell-class specific across brain regions. This confirms the cell-class specificity of baseline firing statistics reported in previous extracellular (Ardid et al. 2015; Constantinidis and Goldman-Rakic 2002; Katai et al. 2010; Mitchell, Sundberg, and Reynolds 2007; Vinck et al. 2013; Tamura et al. 2004) and intracellular (Nowak et al. 2003; Avermann et al. 2012; Druckmann et al. 2013; González-Burgos et al. 2004; Kvitsiani et al. 2013) studies. Furthermore, functionally dissociating four waveform-based cell classes critically extends previous studies that dissociated only two cell-classes based on extracellular recordings (narrow and broad spiking) (Constantinidis and Goldman-Rakic 2002; Mitchell, Sundberg, and Reynolds 2007; Vinck et al. 2013; Thiele et al. 2016; Diester and Nieder 2008). This provides a powerful new window to study cortical circuit function in awake behaving animals.

### Physiological correlates of cell classes

What are the physiological correlates of the four identified cell-classes? With more than two classes, we need to consider subtypes within the excitatory and inhibitory groups. Histological analyses of monkey dlPFC (Krimer et al. 2005), which examined three electrophysiological classes and verified their morphology, showed that broad-spiking RS cells were mostly of the pyramidal type and narrow-spiking FS cells were to a majority GABAergic basket and chandelier cells, as classically described (e.g. McCormick et al. 1985; Nowak et al. 2003). A third intermediate-waveform class consisted exclusively of inhibitory interneurons, with a major proportion of ‘non-fast-spiking’ subtypes (neurogliaform and vertically-oriented cells), which is in line with studies in mice using optogenetic labeling of interneuron subtypes (Avermann et al. 2012; Muñoz, Tremblay, and Rudy 2014; Kvitsiani et al. 2013). The fast-spiking, narrow-waveform profile is typical of parvalbumin-expressing (PV+) interneurons, which morphologically are basket cells. Non-PV+ interneuron types, such as somatostatin-expressing (SOM+) cells, show higher variance in spike width and firing rate, with some overlap with the FS profile. Thus, cell class 1 in the present data (narrowest waveform, high firing rates, low bursting) likely correspond to PV+ fast-spiking interneurons. Class 1 units in LIP also showed phasic visual-evoked responses (Supplementary Figure 2B), consistent with the short time scale of FS units (Kvitsiani et al. 2013) and stronger stimulus modulation described for FS cells (in V4: Mitchell, Sundberg, and Reynolds 2007; in FEF: Katai et al. 2010; Thiele et al. 2016). Non-FS interneurons are likely captured in cell class 2, which shows relatively narrow but more dispersed waveform widths than class 1. The ‘intermediate’ firing rate of class 2 is also in agreement with studies showing differences in firing between FS and non-FS neurons in mice (Avermann et al. 2012; Kvitsiani et al. 2013). The broad-waveform class 4 fits the classical description of RS pyramidal cells, being numerically most abundant in the cortex and having low-rate, regular activity. It is not clear whether class 3 is also part of the excitatory population. A possible clue is given by the relatively strong burstiness specifically of class 3. We can thus speculate that this class comprises intrinsically-bursting (IB) neurons, an electrophysiologically-defined subtype of pyramidal cells that, despite not exhibiting distinct morphology, has often been distinguished from the RS majority based on its atypical firing mode (Connors and Gutnick 1990; Nowak et al. 2003; Katai et al. 2010; Ardid et al. 2015).

The proposed correspondence between the four present classes and physiological cell types is likely to entail some degree of misclassification. For example, some excitatory corticospinal neurons in macaque motor and premotor cortex have FS-like narrow waveforms, with the biggest cells (inferred from axonal conduction velocity) having the thinnest spikes (Vigneswaran, Kraskov, and Lemon 2011). It is not known if this finding applies to other frontal or parietal areas and to what extent this may bias classification. Another case of potential ambiguity between excitatory and inhibitory classes is constituted by ‘chattering cells’, a class of narrow-spiking pyramidal neurons first described in superficial layers of cat visual cortex that can fire high-frequency repetitive bursts in response to stimulation (Gray and McCormick 1996). Although there is some evidence of this cell type in the primate (Friedman-Hill, Maldonado, and Gray 2000; Katai et al. 2010; but see Chen and Fetz 2005), its presence is hard to verify, especially outside of V1 with potentially sub-optimal stimuli as employed in the present study (Gray and McCormick 1996). Complementary morphological, molecular or genetic information (Buzsáki et al. 2015; Zeng and Sanes 2017; Tasic et al. 2018) is needed to unequivocally identify the different physiological cell-types underlying the four cell-classes established here.

### Conclusion

In sum, we show that four functionally distinct neuronal cell classes can be robustly identified from the spike waveform of extracellular recordings across several cortical regions of awake behaving monkeys. These results open a powerful new window to dissect and study the function of cortical micro- and macro-circuits.

## Methods

### Electrophysiological recordings

Extracellular signals were recorded in 70 recording sessions in two rhesus monkeys using Tungsten microelectrodes simultaneously inserted in FEF, dorso-lateral PFC, and LIP. Electrodes were lowered in pairs (1 mm spacing) or triplets (0.7 mm triangular spacing) using custom microdrive assemblies. Electrodes were inserted without targeting of a specific cortical depth. Broad-band extracellular signals were recorded at a sampling rate of 40 kHz and then bandpass-filtered between 0.5–6 kHz to extract spiking activity. The dataset partially overlaps with the multiunit data analyzed in Siegel, Buschman, and Miller (2015).

### Behavioral task

During the recordings, monkeys performed a flexible visuomotor decision-making task. Each trial started with a ‘baseline’ period lasting 0.5 s during which the monkey maintained central fixation. This was followed by a 1 s ‘cue’ period in which a visual cue stimulus was shown to indicate the condition of the upcoming task. Cue stimuli were four different shapes, two of which cued a motion discrimination task and two a color discrimination task. Depending on the cue, the task consisted in judging either the motion direction (up vs down) or color (green vs. red) of a random dot stimulus presented after the cue. The monkeys responded with a leftward or rightward saccade within 3 s after stimulus onset.

### Waveform preprocessing

To obtain spike waveforms, we extracted segments of the filtered voltage traces in a window of 3 ms around each noise threshold-crossing (4 SD; 1ms before crossing) aligned on the main trough of the waveform. The noise level (SD) was robustly estimated as 0.6745 times the median of the absolute of the filtered data. Spike waveforms were trough aligned after spline-based up-sampling. Spike sorting was performed manually offline using Plexon Offline Sorter. Single-unit isolation was assessed by an expert user (CvN) and judged according to a quality index ranging from 1 (clearly distinguishable, putative single unit) to 4 (clearly multi-unit). We used principal components (PC) 1 and 2 of the spike waveform as well as the nonlinear energy function of the spike as axes in 3D sorting space. A putative single unit had to exhibit clear separability of its cluster in this 3D feature space, as well as a clean stack of individual waveforms in its overlay plot. Only units with quality index 1 and 2 were included in the analysis.

We analyzed the average spike waveform of each well-isolated single unit. Waveforms were up-sampled and normalized on its amplitude. We excluded waveforms that satisfied any of three criteria for atypical shape: (1) the amplitude of the main trough was smaller than the subsequent positive peak (n = 41), (2) the trace was noisy, defined as >= 6 local maxima of magnitude >= 0.01 (n = 38), (3) there was one or more local maxima in the period between the main trough and the subsequent peak (n = 35).

### Waveform clustering

As features for cell class classification, we computed two measures of waveform shape: trough-to-peak duration and time for repolarization. Trough-to-peak duration is the distance between the global minimum of the curve and the following local maximum. Time for repolarization is the distance between the late positive peak and the inflection point of the falling branch of the curve (Henze et al. 2000; Nowak et al. 2003).

All preprocessed waveforms (n=2488) were scored on the two measures to obtain a two-dimensional feature space for classification. To identify clusters in the data in an unsupervised way, we used the expectation-maximization (EM) algorithm for Gaussian mixture model (GMM) clustering. We modelled the data as a weighted sum of multivariate Gaussians:

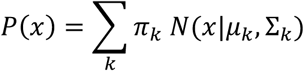

with *k* components parametrized by mean *μ*, covariance Σ and mixing coefficient *π*. The EM algorithm fits this model by iteration of a two-step process: it first estimates posterior probabilities of the data given the current set of parameters (E step), and then updates the parameters to maximize the log-likelihood function of the model given the current estimates (M step). The steps are repeated until convergence. We initialized the process with random parameters for 50 repetitions and chose the fit with the largest log-likelihood among the replicates.

To select the number of Gaussian components in the model we used the Bayesian information criterion (BIC) (Schwarz 1978):

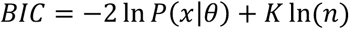

where *P*(*x|θ*) is the maximized likelihood for the estimated model, *K* is the number of parameters, and *n* is the sample size. By including a penalty term that grows with the number of parameters, the BIC cost function effectively favors simpler models and reduces overfitting. The optimal number of clusters was chosen as the value that minimized the BIC computed between 2 and 10 components.

After fitting the model, we determined cluster memberships by ‘hard’ assignment: each unit was assigned to the class associated with the highest posterior probability.

Given the initial clustering outcome, we excluded one high-variance cluster that captured the waveforms dispersed around the high-density axis of the data cloud (n = 212). We also excluded units that were outliers of the whole data cloud, defined as having Mahalanobis distance large than 5 from the centroid of the Gaussian cluster they were assigned to (n = 69). After outlier rejection, we re-ran the clustering (including the BIC analysis for choosing number of components) to obtain the final cell class classification.

To assess the degree of cluster separation, we calculated the overlap between GMM components using a Monte Carlo approach. We randomly generated 10000 data points from the fitted GM distribution and compared, for each data point, the true cluster from which the observation was drawn with the class to which it was assigned. The outcome of this comparison can be represented by a confusion matrix, where the off-diagonal terms provide empirical estimates of the area of overlap between the GMM components. Overall class separation was quantified as the mean of the diagonal probabilities of the confusion matrix.

This method was extended to assess the similarity of the clustering schemes obtained for individual cortical areas. For each pair of areas A and B, 10000 data points were randomly drawn from the GM distribution of area A and assigned to classes defined on the GM distribution of area B. The confusion matrix shows, for area A’s data, the true generating cluster against the assigned class label. For all area pairs, overall class similarity was quantified as the mean of the diagonal probabilities of the corresponding confusion matrix.

### Analysis of firing statistics

To characterize spontaneous activity, we analyzed spiking activity during the baseline fixation period. We averaged across baseline periods of all trials. We computed four firing statistics: mean firing rate across trials (FR), Fano factor (variance over mean of spike counts across trials, FF), coefficient of variation of the inter-spike interval distribution (CV_ISI_) and burst index (BI). Both Fano factor and CV_ISI_ are mean-standardized measures of dispersion that reflect firing regularity, with an expected value of 1 for Poisson firing and values below 1 indicating more regular firing (Shinomoto, Shima, and Tanji 2003). The burst index was defined as the ratio between the observed proportion of bursts, defined as inter-spike intervals < 5 ms, and the proportion of bursts expected from a Poisson process with equal mean rate (Constantinidis and Goldman-Rakic 2002). This measure quantifies the tendency to fire in bursts unbiased by firing rate. To avoid under-sampling, we only computed the burst index for units with more than 50 inter-spike intervals pooled across all trials (n=1388 units), which excluded neurons with very low firing rates.

We tested for significant differences in activity between cell classes using a one-way ANOVA on each firing statistic. All measures were log-transformed to optimize normality. To control for the unequal regional distribution of cell classes, we randomly subsampled the data to have equal cell class proportions in the three areas (‘area-stratified datasets’). The ANOVA F-statistic was computed as the ratio of mean square between and mean squared error. Both numerator and denominator were calculated on 1000 area-stratified subsamples and then averaged across subsamples, so that the F-ratio was obtained from the two averaged quantities. For post-hoc comparisons, we computed pairwise t-tests using the average difference of means and the average pooled standard error across 1000 area-stratified subsamples. We corrected for multiple tests using False Discovery Rate (FDR) (Benjamini and Hochberg 1995) correction.

### Multivariate decoding

We performed cell class decoding using Support Vector Machine (SVM) classification on several different feature sets. For all feature sets, decoding was performed on 50 area-stratified subsamples (where cell class proportions were matched across cortical regions by random subsampling) and averaged across subsamples to obtain the final classification estimate.

#### Classification procedure

To reduce the multiclass problem to binary classification, we independently trained and tested six binary SVMs for each pair of cell classes. The six sets of predicted labels were combined by majority vote (‘one-vs-one’ classification); in case of ties, one of the two winning classes was chosen at random. The SVM algorithm employed a Gaussian radial basis function kernel with a scaling factor of 1.

Each binary classifier was evaluated using 10-fold cross-validation. Within each classifier, we randomly subsampled the data such that both cell classes had N equal to the minimum sample size across all cell classes. Equal cell class proportions were preserved in each cross-validation fold’s training and test set, so that chance-level classification performance for a given test set was always 0.5. By using the global minimum of sample sizes across cell classes, we also ensured that all pairwise classifiers had comparable signal-to-noise ratio. This stratification procedure was repeated 100 times and the estimates were combined by majority vote.

Classification outcome was summarized in a confusion matrix of probabilities based on the average counts over 50 area-stratified datasets. Counts were divided by true class total counts to obtain the probability of predicting each class given the true label. Each class was considered to be decodable if its true positive rate was significantly greater than chance level of 0.25 in a binomial test, corrected for four tests using FDR correction. As a summary measure of the confusion matrix, we quantified classifier accuracy as the average true positive rate across cell classes (mean of the diagonal of the confusion matrix).

#### Cross-area classification

To assess area specificity of cell class decoding, we trained classifiers on data from one cortical area and used them to predict data from other areas. We matched areas’ signal-to-noise ratio by creating 50 randomly subsampled datasets for which all areas had the same number of observations, equal to the minimum N across the three areas. A separate classification instance was run for each subsample, and the resulting 50 confusion matrices of counts were averaged.

To test for significance of cross-area classification performance, we used a permutation test that compared the observed accuracy with an empirical null distribution. The null distribution was constructed by training classifiers with randomly permuted class labels. We ran 1000 instances of null classifiers, each resulting from the average across 20 area-stratified subsamples. The observed accuracy was declared significant if it exceeded the accuracy values of 1000 label-shuffled classifiers at p<0.05, FDR-corrected for 9 tests (3*3 cross-area classifiers).

Area specificity was computed from the 3-by-3 matrix of cross-area classification accuracy as the ratio between the average of the diagonal and the average of the non-diagonal values.

### Linear Discriminant Analysis for feature importance estimation

For the dataset comprising all features (four baseline firing statistics, PSTH, cue information) we also trained a Linear Discriminant Analysis (LDA) classifier using the same procedure described above for the SVMs. To quantify each feature’s contribution to the decoding, we computed FIRM (Feature Importance Ranking Measure) using the simplified formulation for unregularized OLS regression (Zien et al. 2009; Haufe et al. 2014):

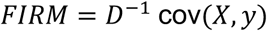

where *D* is a diagonal matrix of standard deviations of the features, *X* is the training data matrix, and *y* is the vector of true class labels of the training data (recoded as +1 and −1). As discussed in Haufe et al. (2014), FIRM for linear decoding can be approximated by the covariance between the data and the labels, effectively reducing to a univariate analysis on each feature. The normalization by *D* ensures that FIRM is invariant to feature scaling. As our measure for feature importance we considered the magnitude (absolute value) of FIRM.

### Principal component decomposition of PSTH

Peristimulus time histograms (PSTH) of single-unit spike counts were computed using 50 ms bins, within a 1.5 s trial window comprising the 0.5 s baseline fixation period and the 1 s cue period. Each unit’s PSTH was z-scored on the mean and standard deviation of the baseline period across trials.

We extracted low-dimensional PSTH features using Principal Component Analysis (PCA). The number of significant principal components was determined using cross-validation (Bro et al. 2008). For each number of PCs *k*, we fit a PCA to one portion of the data (‘training’ set) and, using the first *k* components, we reconstructed the left-out data (‘test’ set). Importantly, to obtain reconstructions that were truly independent from the data, we excluded in turn each variable from the fitting on the training set, and that variable only was predicted for the test set observations. Using 10-fold cross-validation, a predicted value was calculated for each data point. The overall prediction error was calculated as the sum of squared differences between true and reconstructed data points (PRESS, Prediction Residual Error Square Sum) as a function of number of components:

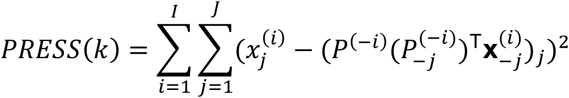

with *k* number of PCs, data samples *x*_*ij*_ having number of observations *I* and number of variables *J*, and *P* being the *J* × *J* matrix of PCA coefficients, with columns corresponding to principal components. The number of significant components was chosen as the global minimum of the PRESS curve.

PCA feature extraction, including selection of number of PCs, was performed independently on each of the 50 area-stratified datasets for cell class classification. For cross-area classification, the PCA transformation was estimated on the training area only and then applied to data of the test area.

### Cue information

Cue information was quantified as the effect size of the cue-factor in a 7-way ANOVA computed on the late cue period of the trial (500–1000 ms after cue onset) (Siegel, Buschman, and Miller 2015). Effect size was quantified by the ω^2^ statistic, an unbiased estimator of the variance explained by an ANOVA factor independently from all others factors in the model (Olejnik and Algina 2003):

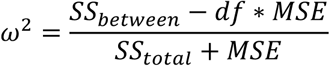

where *SS*_*between*_ is the between-groups sum of squares, *df* is the degrees of freedom, *SS*_*total*_ is the total sum of squares, and *MSE* is the mean squared error. The first three factors of the ANOVA corresponded to the cue of each trial grouped into two levels according to all three possible pairwise pairings of the four task cues. We computed cue information as the average explained variance of all first three factors (Siegel, Buschman, and Miller 2015). The remaining 4 factors of the ANOVA were the motion direction of the random-dot stimulus, the color of the random-dot stimulus, the response of the animal on the current trial, and the response on the previous trial.

To control for linear dependencies between cue information and activity measures, we took the residuals of cue information after regressing out the four baseline firing statistics (firing rate, Fano factor, coefficient of variation of the ISI distribution, burst index).

Cell class differences in cue information were assessed with a one-way ANOVA using the same procedure employed for the firing statistics: the data was subsampled 1000 times to match cell class proportions across cortical areas, and the F-ratio was calculated from the averages of the numerator and denominator across subsamples. Post-hoc pairwise comparisons between classes were similarly evaluated taking the average numerator and denominator of the t-statistic.

## Acknowledgements

This work was supported by grant NIMH 5R37MH087027 (E.K.M), the European Research Council (ERC) StG335880 (M.S), the Deutsche Forschungsgemeinschaft (DFG, German Research Foundation) project 276693517 (SFB 1233) (M.S.) and grant SI1332-3/1 (M.S.), and the Centre for Integrative Neuroscience (DFG, EXC 307) (M.S.).

## Author contributions

Conceptualization: M.S., C.T., E.K.M; Methodology: C.T., M.S., C.v.N; Investigation: M.S.; Formal Analysis, C.T., M.S., C.v.N; Writing – Original Draft: C.T., M.S.; Writing – Review & Editing: C.v.N; Funding Acquisition: M.S., E.K.M.; Resources: M.S., E.K.M.; Supervision, M.S.

## Supplementary Figures

**Supplementary Figure 1.**
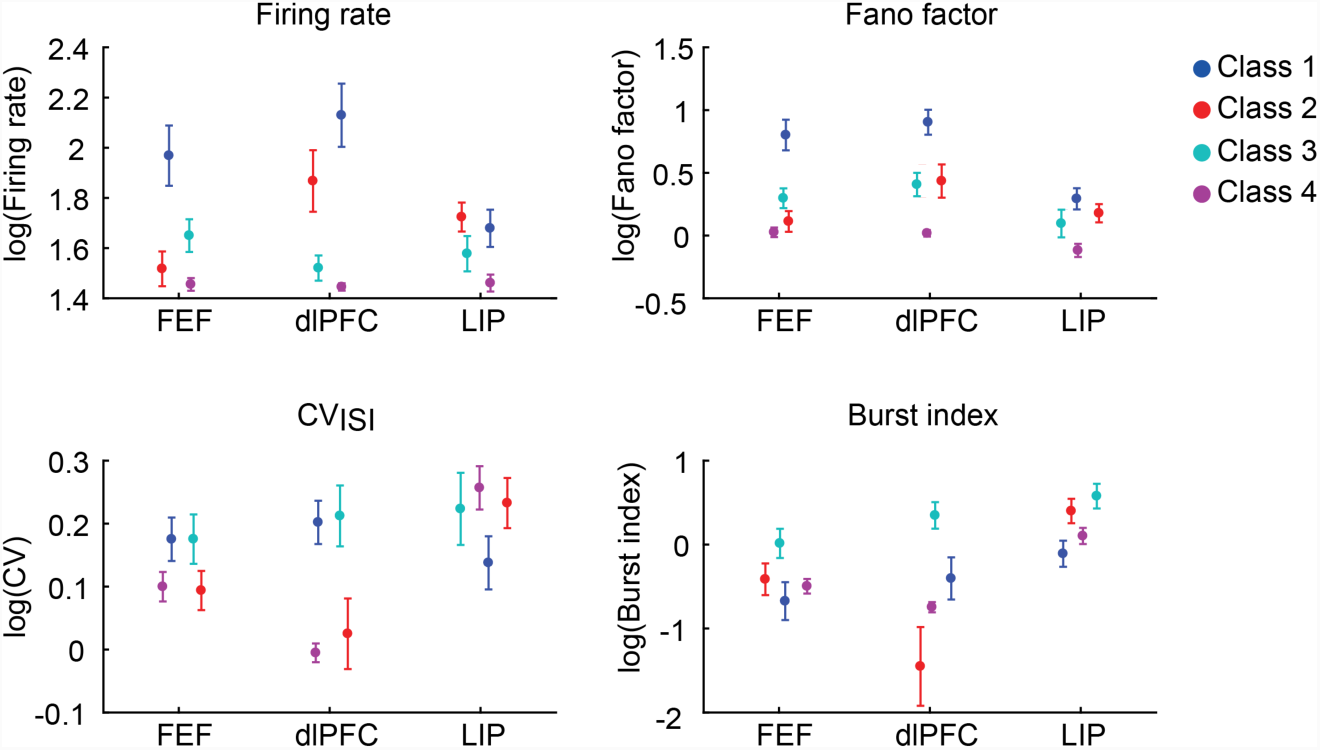
Baseline firing statistics by cell class and cortical area. Firing statistics by cell class, separately for each area. Error bars denote SEM.

**Supplementary Figure 2.**
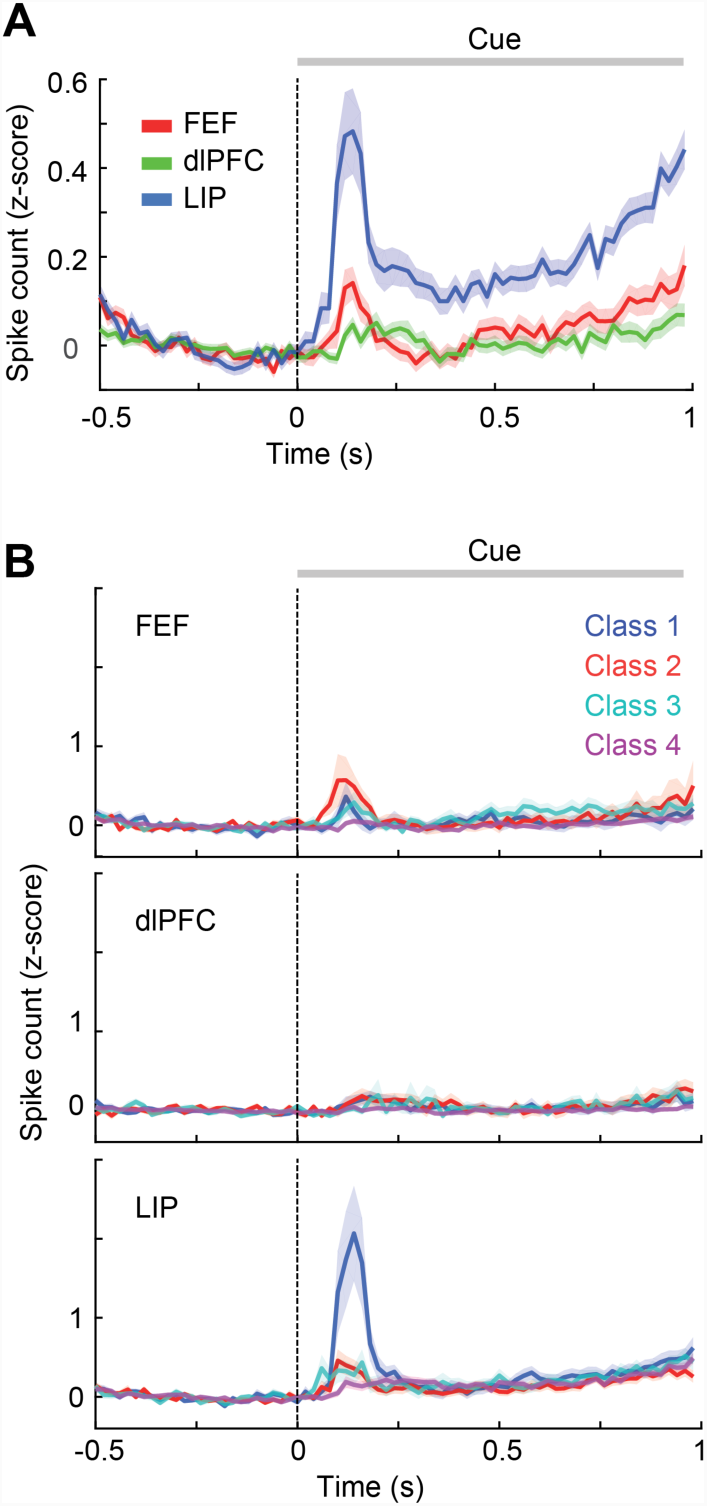
Response dynamics by cortical area. (A) Average PSTHs for single-units recorded in the three areas FEF, dlPFC and LIP. PSTHs were computed in a 1.5 s window comprising the baseline and cue period of each trial (20 ms bins for visualization purposes; all decoding analyses used 50 ms bins). Error bars denote SEM across units. (B) PSTHs by cell class within each cortical area.

**Supplementary Figure 3.**
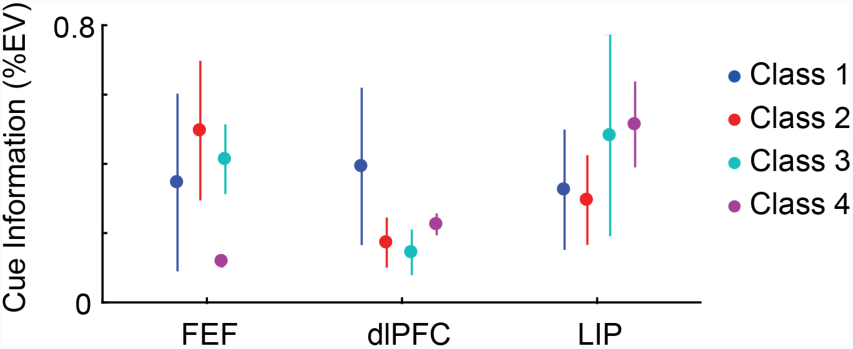
Cue information by cortical area. Cell class means and SEM of cue information for each cortical area.

